# Isolation of individual natural killer cells from deep, high-aspect ratio microwell arrays - an evaluative study

**DOI:** 10.1101/2025.02.20.639369

**Authors:** Quentin Verron, Niklas Sandström, Hanna van Ooijen, Karolin Guldevall, Karl Olofsson, Thomas Frisk, Björn Önfelt

## Abstract

Immune cells exhibit functional heterogeneity beyond what is resolved by classical definitions of subpopulations based on cell surface expression of receptors. To develop efficient and personalized cell-based immunotherapies, we need to resolve this heterogeneity and understand the underlying parameters that dictate cellular responses to specific target cells. For this, new methods are required that can identify and harvest immune cells with specific functions, e.g., high cytotoxic potential, to form clonally expanded cells or to assess molecular or genetic signatures. In this study, we evaluate a system for non-destructive, live cell picking and release in deep, high-aspect ratio microwells and test it for isolation of individual natural killer (NK) cells. We assess its performance at retrieving and releasing beads from microwells and demonstrate its potential for single NK cell isolation with intact viability. We also implement a semi-automated workflow for functional single-cell screening of NK cell behavior in microwell arrays followed by single-cell identification and isolation, demonstrating the potential for functional screening and isolation of serial killing immune cells. Our evaluation concludes that this cell isolation system, in combination with microwell arrays, offers opportunities for improved understanding of NK cell biology with applications towards cell therapy. However, its limited throughput hinders large-scale applicability.

## Introduction

Natural Killer (NK) cells are cytotoxic lymphocytes known for their ability to recognize virus-infected and transformed cells, including tumor cells. This surveillance is orchestrated by activating and inhibitory signals mediated by the ligation of a range of receptors at the NK cell surface with their cognate ligands on the target cell (1). The expression of these receptors varies from cell to cell, resulting in a vast pool of phenotypically different cells (2). The receptor repertoire has historically been used to discriminate between NK cell subpopulations, with some phenotypes that can be correlated to key functional attributes, such as activation, education and cell maturation (3–6). Recently, the analysis of gene and protein expression, complemented with other omics outputs, have in many instances replaced traditional analysis of surface receptor expression for more detailed, multi-dimensional segregation of phenotypically distinct NK cell subsets (2,7–9). One common trait to many of these new approaches is that possible functional differences between sub-populations can be hypothesized *a posteriori*, but not easily verified.

True *a priori* identification of functionally distinct cell subsets is by nature a challenging process. Extensive functional interrogation of heterogeneous populations must be performed over thousands of cells with single cell resolution and complemented with proteomic or genetic characterization of the cells identified by their functional properties. Conventional experimental approaches for functional studies are often not based on single cell readouts and rarely offer a link to downstream analysis. Flow and mass cytometry offer single-cell resolution with high throughput but with limited possibilities for direct functional readouts (10–13). Imaging flow cytometry (14,15) offers spatial information of molecular distribution in individual cells at high-throughput, which can be used as the basis for cell sorting or phenotypic characterization. However, the imaging information is based on a single time point, limiting the applicability of the method for dynamic functional readouts. Droplet-based microfluidic systems (16–19), also provide high-throughput with the possibility to follow cellular behavior or to perform various forms of genetic or molecular analysis. However, with a few exceptions, there is rarely a link between the two forms of analysis (20). Other microfluidic systems for functional screening and single cell retrieval have been demonstrated, but they often require complex and sensitive fluidic setups that are specialized for, and limited to, only one type of functional screening (21,22).

We and others have used microscopy-based live cell screening in microwell arrays as an alternative to flow-based systems (23–31). By confining cells in spatially separated microwells, longitudinal functional studies of individual cells can be performed over periods of several days. Such systems have been used to identify populations of highly cytotoxic NK cells, so-called serial-killing NK cells (25,28,32–34), which exhibit a specific behavior in terms of contact dynamics and delivery of lytic hits towards tumor cells (33,35). As NK cells of interest remain confined in their microwell after a screen, there is potential in retrieving identified cells for further downstream analysis. Various isolation methods have been implemented for single cell retrieval from arrays of deep microwells, but often at the cost of the destruction of the culture vessel (36–40). In other systems, single-cell isolation has been achieved using precisely controlled micropipettes whereby cells are aspirated directly from the microwells (23,24,31,41– 43). However, these microwell arrays were often made of soft polymers, and had microwells of a shallow depth, which increases the risk of cells escaping the microwells during media exchange and long-term assays.

In this work, we evaluate and demonstrate how imaging-based functional screening of NK cells in deep microwells can be combined with individual isolation of cells of interest using a piezo-actuated picolitre micropipette (CellSorter Scientific). We characterize the cell isolation performance and demonstrate the ability of the system to specifically transfer individual cells between microwells with retained viability. In addition, we illustrate screening based on cytotoxicity of primary NK cells against leukemia cells, followed by successful isolation of serial-killing NK cells. Together, our results confirm the potential of this semi-automated system for functional cell screening combined with specific single cell isolation. However, the limited throughput of the method restricts the analysis to downstream investigations compatible with small cell numbers.

## Results

### Workflow for isolation of single cells after functional prescreening

To achieve single-cell isolation in microwells, we used a top-mounted, micropipette-based isolation system integrated with an inverted wide-field light microscope (Fig 1A). The isolation system is semi-automated and controlled by computer software. Importantly, it fits within the environment control chamber of the microscope, making it compatible with long term assays in a closed and contamination-free environment. The micropipette head includes a LED-based light source for providing transmission light from above the chip, in addition to the standard excitation light for fluorescence of the inverted microscope.

**Figure 1:**
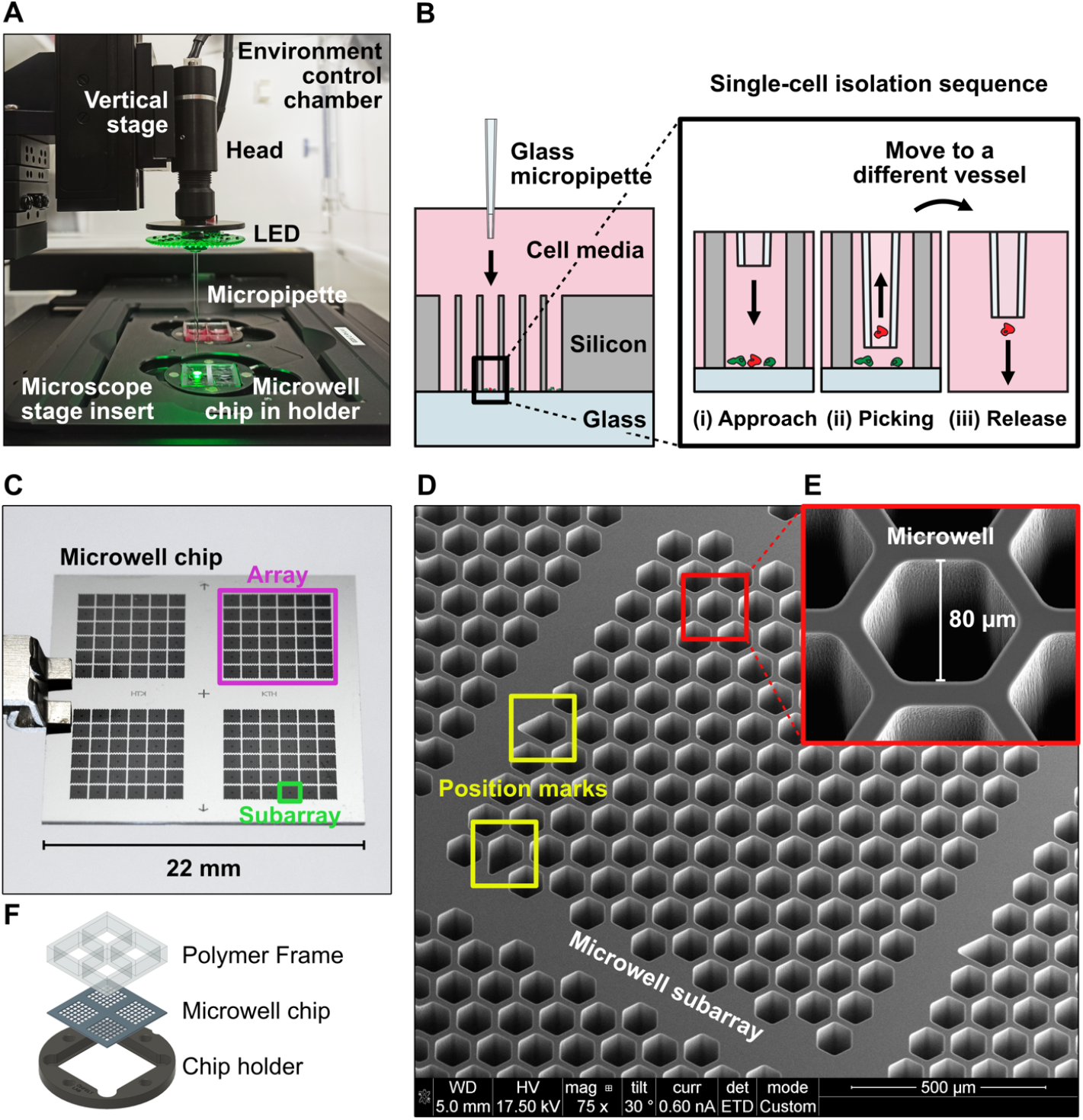
Single-cell isolation in deep microwells. **(A)** Photograph of the integrated setup for functional screening and single-cell isolation. The cell isolation device is installed on an inverted fluorescence microscope equipped with a motorized stage and a chamber for environmental control. The microwell chip is mounted in a custom-made holder, which in turn is placed in a commercial stage insert for 35 mm tissue culture dishes. **(B)** Working principle of single-cell isolation from the microwells. A thin glass micropipette is used to access the wells from above. Three critical moments are illustrated: (i) Approach, as the micropipette is lowered to 15 µm directly above the cell of interest; (ii) Picking, whereby the cell is gently aspirated into the micropipette opening; (iii) Release, when the cell is deposited into a different culture or analysis vessel. **(C)** Photograph of a silicon-glass microwell chip with four separated well arrays, each including 36 subarrays. **(D)** Electron micrograph of a microwell subarray, including 143 microwells in a honeycomb layout and corresponding to the size of an imaging FoV with a 10x magnification objective. **(E)** Electron micrograph of a hexagonal-shaped microwell. **(F)** Illustration of an exploded view of the four-chamber polymer frame, which is bonded to the Silicon-glass microwell chip, and the bonded chip is placed in a custom-made chip holder in a 35 mm tissue-culture dish format.

One critical aspect of the cell isolation system is its compatibility with the microscale format of the deep and narrow microwells used in our silicon-glass microchip (Fig 1B). The system depends on a piezo-electric actuator for precise fluid control of both volume and flow rate for gentle handling of the isolated cells. The glass micropipette has an outer diameter that is less than the width of the microwells and a length that is greater than the depth of the microwells. Thus, the micropipette reaches all the way down to the well floor and it can be laterally positioned to target specific cells in the wells. The lateral movement of the micropipette in relation to the microwell array is performed by the microscope stage, whereas the vertical movement is controlled by a separate single-axis stage controller of the micropipette head, both of which feature micrometer precision. In this way, the micropipette can be aligned to specific microwells on the chip and then be lowered into a well to a set distance above the cell of interest. The isolation procedure includes three steps (Fig 1B): (i) approaching the cell of interest in the microwell; (ii) picking up the cell from the microwell through aspiration; and (iii) releasing the cell into another microwell or other culture vessel by dispensing. Cell aspiration and dispensing can be monitored by imaging using either transmitted light or fluorescence (Supp movie 1). If multiple cells are of interest, they are isolated in a sequence. Thereafter, the isolated cells can ideally be further characterized, expanded into larger numbers or used in other ways depending on the application.

In this work, a custom Matlab-based software for analysis of imaging data from NK functional screening assays in the microwell chips (44) has been adapted for bridging NK screening assay data curation with single-cell isolation. The microwells are directly selected for cell isolation according to the results of the functional assay performed in the chip. The microwell array positions of the wells of interest are converted to microscope stage coordinates and transferred to the proprietary software controlling the micropipette isolation system. One key advantage of using the microchip platform compared to conventional cell culture substrates is that the layout and design of the well array makes it simple to identify the positions of individual microwells and cells even if different microscopes are used for analysis and cell retrieval.

We have thus designed a workflow combining functional screening of NK cells in the microwell array chip, allowing identification of individual NK cells with a certain functional response and subsequent specific retrieval of these cells from the microwells for release into a separate microwell or other culture vessel for downstream analyses. Some key components of the workflow are evaluated individually before being combined in a proof-of-principle experiment, as shown in the figures that follow.

### Design of microwell array chips for functional screening assays and subsequent cell isolation

The microwell array chips used in this work have a design that is optimized for both functional screening assays and single-cell isolation with the micropipette system. The chip includes four microwell arrays (Fig 1C), each consisting of 36 subarrays (Fig 1D), which correspond to the area of the imaging field-of-view (FoV) using a 10x magnification objective on the microscope used for screening. During screening, the microscope scans the entire chip by imaging the 144 subarrays. Each subarray includes two irregularly shaped microwells that function as position marks, making it possible to trace back the position of each microwell from the imaging data.

The microwells have a hexagonal shape and are distributed in a honeycomb layout, maximizing the fill factor of wells in each array. The hexagonal shape of the microwells enables full compatibility with the use of the circular glass micropipette, allowing easy access and movement inside the wells. The smallest width of the microwells is 80 µm and the depth is 300 µm (Fig 1E). The aspect ratio and the small size of the wells permits functional screening assays of single NK cells, and the cells are confined without risk of escaping during long periods of time. Each subarray, i.e., microscopy FoV, contains 143 microwells. In total, the 22×22 mm^2^ chip includes 20 592 microwells.

The microwell chip is made of solid, high-quality materials; the micro-structured top layer, which includes the wells, is produced in silicon, while the bottom layer is made of 175 µm thick glass, with similar properties to a #1.5H glass coverslip. Such a design allows for high-resolution imaging with inverted microscopes. Upon use, the chip is placed in a custom-made holder in the shape of a cell culture dish that fits directly in standard microscopy stage inserts (Fig 1F). A four-chamber polymer frame is bonded on top of the chip so that the chambers align with the four microwell arrays on the chip. Each chamber is accessible by manual pipetting from above, and is used for adding and replacing liquids, such as cell suspensions and cell culture media. Thus, four different experimental conditions can be used simultaneously during a functional screening assay.

### Characterization of isolation performance in microwells

As the piezo-actuated isolation system is designed for retrieval of cells from open culture vessels such as Petri dishes or shallow well arrays, we decided to characterize its isolation performance in combination with the specific geometry of our microwells. For this purpose, we used NK cell-sized silica beads as a model. After priming the micropipette and actuation chamber with liquid, we verified the performance of the system ahead of each experiment (Fig 2A). Beads were seeded in a Petri dish and 10 beads were randomly selected for isolation. The test was considered “passed” if over 50% of the target beads were successfully picked and released. We identified the picking phase as the most sensitive of the isolation process, with 81.8% successful picking events during passed calibrations (Fig 2B) compared to 94.5% successful release events (Fig 2C). We did not observe any clear difference in performance across these ten successive retrieval events, suggesting a good stability of the system during isolation sequences (Fig 2B-C).

**Figure 2:**
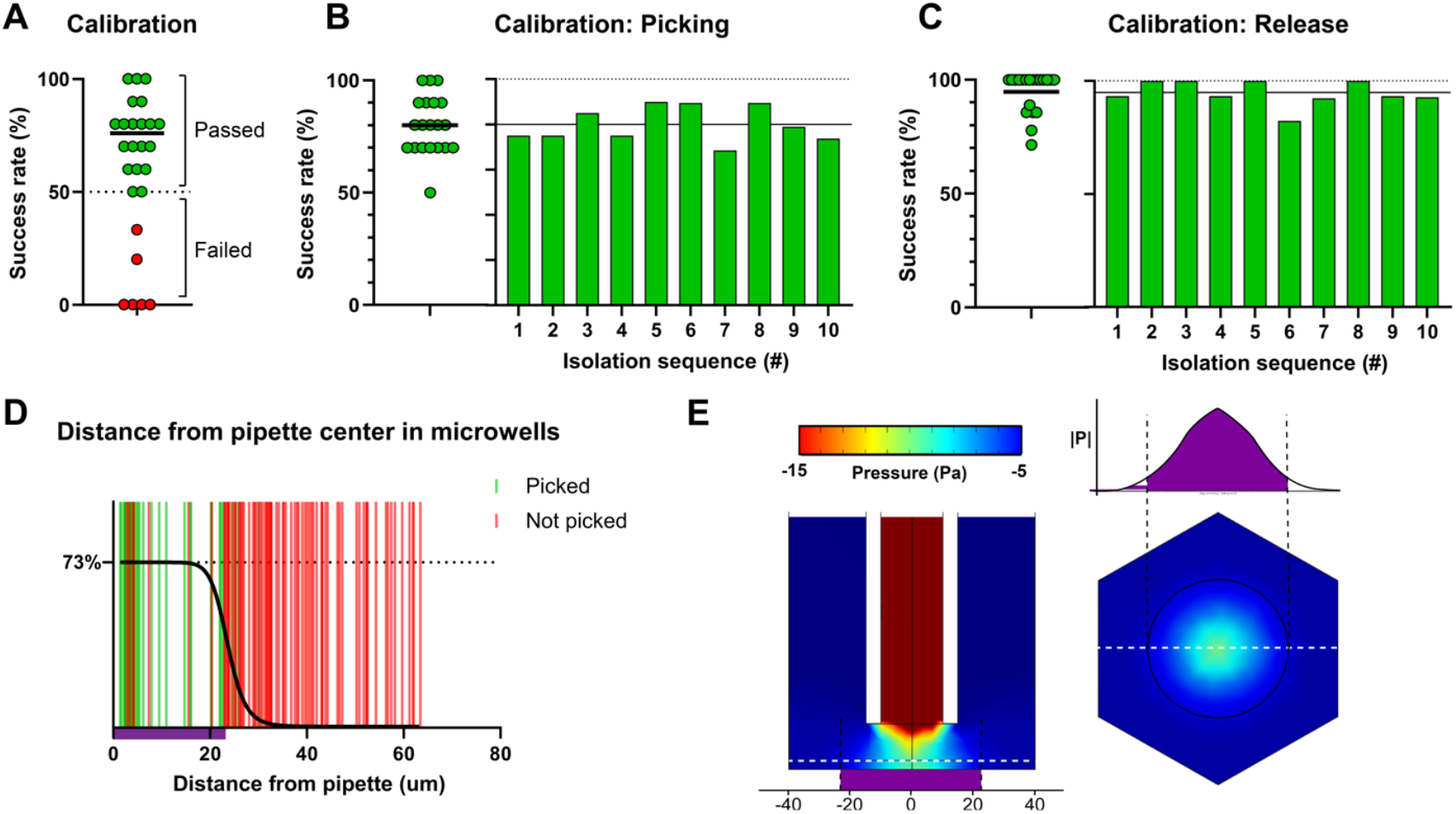
Characterization of the isolation performance. **(A-C)** Calibration of the isolation system with silica microbeads in a cell culture dish with the pipette placed 15 µm above the beads. **(A)** Average success rate of isolation of 10 randomly selected beads. The calibration is considered passed if over 50% of the beads are successfully retrieved. The black line represents the overall mean value of the successful calibrations. **(B)** Success rate of picking during passed calibrations, plotted as an average of each 10-bead sequence (left), or as a function of the number of the bead in the picking sequence (right). **(C)** Success rate of release during passed calibrations, plotted as in (B). **(D)** Picking outcome as a function of the lateral (x, y) distance of the bead from the pipette center. The analysis included both the targeted beads (distance to the pipette center below 10 µm) and neighboring beads. The radius of effect (indicated in purple) and probability of success were estimated by fitting a four-parameter logistic curve (black line). **(E)** Finite element model simulations of fluid pressure for a 20 µm inner diameter micropipette at 15µm from the bottom of a hexagonal 80 µm-wide microwell. Left: side view, with the horizontal section at 3µm from the bottom indicated by a white dashed line. Right: horizontal section, with the experimentally measured area-of-effect in overlay, and the calculated pressure along the white dashed line plotted above.

Moving into the microchip, we also used silica beads to characterize the isolation from our wells. To understand how picking is affected by the geometry of the wells, we evaluated the probability of a bead getting picked depending on its lateral distance from the pipette center (Fig 2D). We found that all beads successfully picked lay within 30 μm laterally from the center of the pipette, a result consistent with the 30 μm-sorting resolution of cells in Petri dishes measured by the manufacturer (45). The distance-dependent picking probability was estimated by fitting a four-parameter logistic curve on the outcome data (with 1 being a picked bead and 0 a bead not picked). We obtained a maximum picking success rate of 73% closest to the pipette, with half that probability reached within 23 μm, which we thus define as the radius-of-effect of the pipette (Fig 2D). Indeed, in our experiments, some beads were observed to be displaced by the micropipette aspiration but not picked, and these were found on average 26 μm from the center of the micropipette i.e., at the edge of the area-of-effect (Supp fig 1A). The radius-of-effect that we measured was consistent with the results from the corresponding fluid flow simulations, as we found that the negative pressure created by the micropipette diminished just beyond that radius (Fig 2E). When two beads were found within 23 μm from the pipette center, we measured that the probability to pick either bead, both or none, was as predicted by assuming that picking each bead is an independent event with a 73% probability of success (Supp fig 1B). In other terms, the micropipette displayed a sorting resolution of 23 μm when aiming at the center of the beads. As flow simulations predict the flow profile around the micropipette to be markedly altered close to the microwell walls (Supp fig 1C-E), we investigated whether unaffected beads within the radius-of-effect of the pipette laid closer to the wall. However, we found that at a matching distance from the pipette center, there was no measurable effect of the distance to the wall (Supp fig 1F). Overall, our results confirm that the piezo-actuated isolation system can be used to successfully pick beads from our silicon-glass microwells, with a 23μm resolution and 73% success rate.

### Isolation of primary human NK cells

We investigated the performance of the system to isolate primary NK cells from the microwells. Single NK cells were seeded in the microwells, then cells found farther than 15 μm from the well wall were randomly selected for isolation. The NK cells were successfully retrieved from the wells and released in another chip compartment, albeit with a lower success rate than achieved using silica beads (Fig 3A). To investigate whether this was due to cells adhering to the chip or micropipette walls, we compared normal culture medium to medium supplemented with EDTA to decrease cell adhesion (Fig 3A-C). We found no difference in the overall success rate, or the picking or release phases with the two different culture media (Fig 3B-C). Similar to the results obtained with calibration beads, we observed no significant difference in performance for cells isolated successively in a sequence (Fig 3B-C).

**Figure 3:**
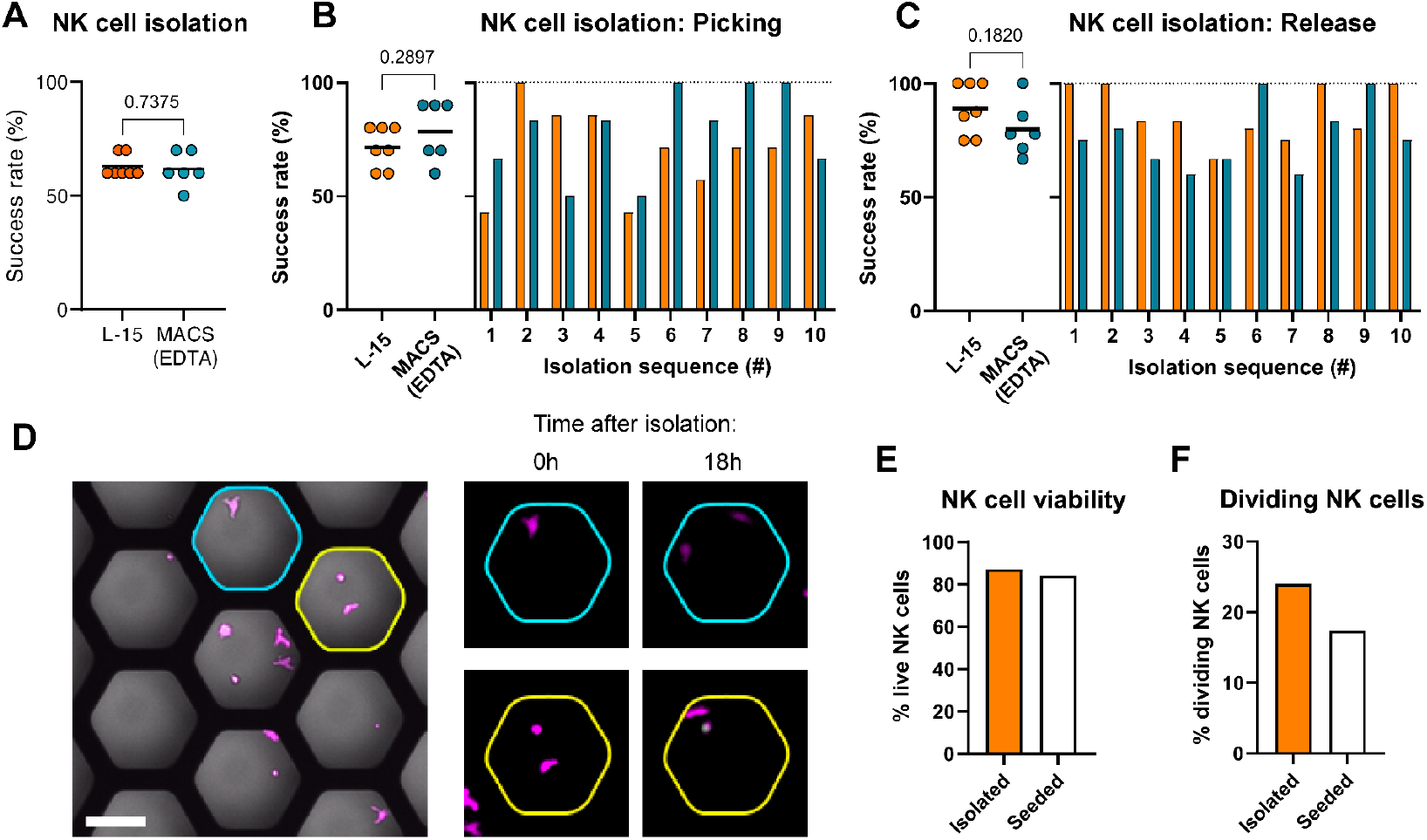
Isolation of single NK cells from microwells. **(A-C)** Success rate of isolation of primary human NK cells incubated in microwells containing either L-15 or magnetic separation medium (MACS), divided into overall success rate (A), success rate of picking, plotted as an average of each 10-cell sequence (left), or as a function of the number of the cell in the picking sequence (right)(B), or success rate of release (C), plotted as in (B). **(D)** Primary NK cells incubated in microwells were identified by imaging (left), picked and released into individual microwells on a different microchip, and monitored for 18 hours by time-lapse imaging (right). Top row (cyan): The isolated NK cell divided during the time-lapse sequence. Bottom row (yellow): One of the two isolated NK cells died during the time-lapse sequence. Scale bar: 50 µm. **(E)** Viability of the primary NK cells 18 hours after isolation, compared to cells manually seeded at matching density. (**F**) Fraction of cells dividing within 18 hours after isolation. (E-F): Isolated: N=25, seeded: N=75.

It is often desirable to retrieve live cells from the microwells, either for expansion or for subsequent experiments. Therefore, the effect of the isolation process on the viability of primary NK cells was investigated. We isolated individual primary NK cells from the microwells in one compartment of the chip and released them in wells in a different compartment, then monitored them by time-lapse imaging for 18 hours (Fig 3D). We found that the isolated NK cells had a comparable viability to manually seeded cells (Fig 3E), with some cells dividing within the first 18 hours after isolation (Fig 3F). Our results confirm the potential of our workflow for use with sensitive suspension cells such as primary NK cells, with a significant fraction of the cells remaining viable after the isolation process.

### Identification and isolation of serial-killing NK cells

Having confirmed that our setup could support single-cell retrieval from the microwells, we investigated whether it could be combined with microchip screening to achieve isolation of specific cells based on functional readouts (Fig 4A). For this purpose, we conducted a cytotoxicity screen by challenging primary human NK cells with K562 leukemia target cells (Fig 4B-C). At the end of the assay, the chip was moved to an incubator while we proceeded to analyze the data. As previously described (33), we observed that a minority of NK cells performed multiple kills in sequence (Fig 4D-G). After identifying these serial killer NK cells, the chip was transferred to the microscope dedicated to single-cell isolation, and the microwell array coordinates were used to locate the wells of interest (Fig 4H, image i.). The NK cells could be successfully picked (Fig 4H, image ii) and released into a microwell of a different compartment of the chip (Fig 4H, image iii). The removal of the cells of interest from their source well was confirmed in a post-isolation microscopy scan of the chip (Fig 4H, image iv). These results show the potential of our setup for on-chip single-cell screening where cells are identified based on functional readouts followed by specific retrieval of pinpointed cells (Supp movie 1).

**Figure 4:**
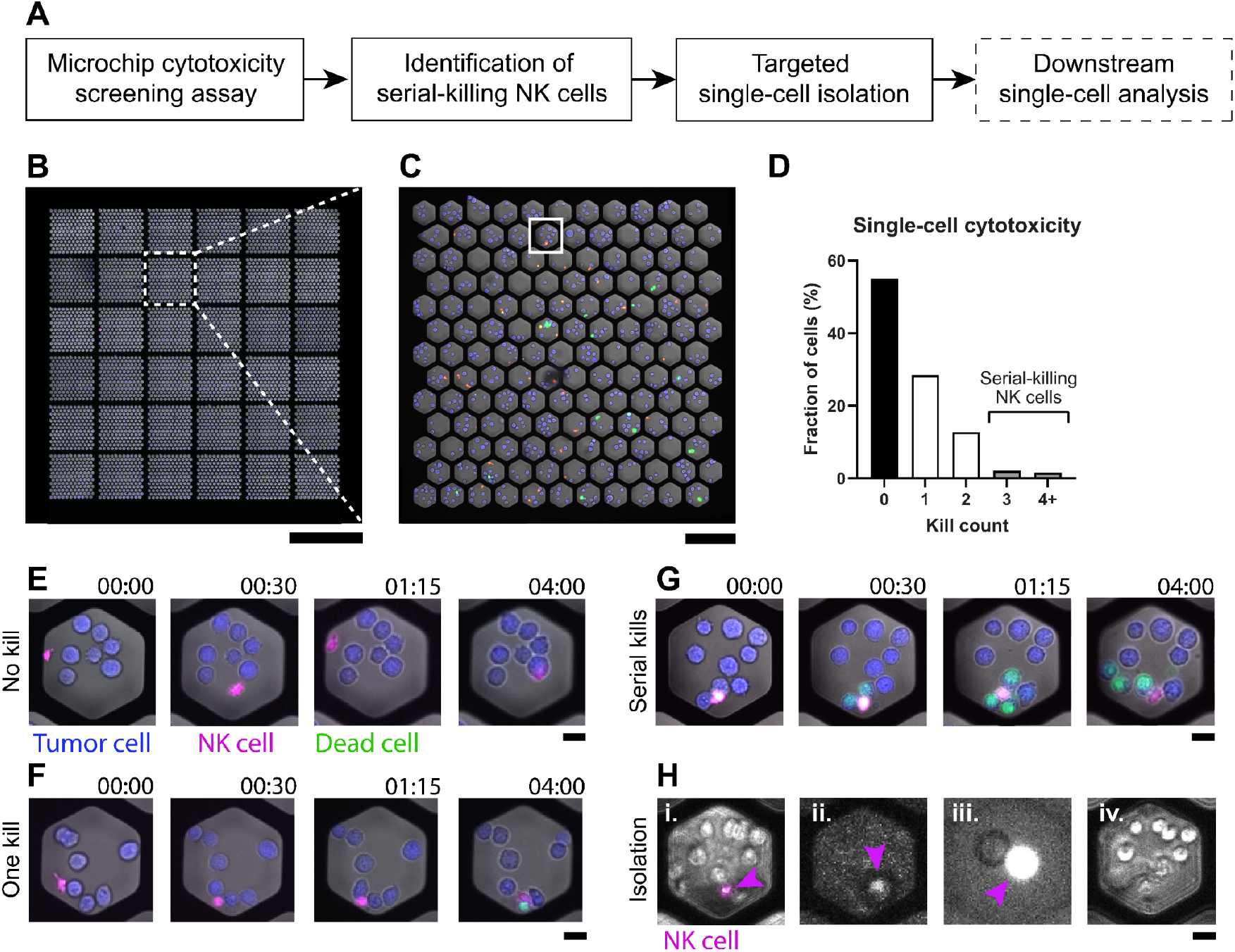
Isolation of individual NK serial killers identified from cytotoxicity screening assay. **(A)** Flowchart of selective isolation of NK cells based upon the results of a cytotoxicity screening assay. **(B-D)** Primary human NK cells and K562 tumor target cells were distributed on the microwell chip at an E:T ratio of 1:5, at a concentration that maximized the number of wells with individual NK cells. **(B**) Overview of the microchip assay showing one microchip chamber. **(C)** A single imaging FoV containing 143 wells. The orange outline indicates a well containing a serial-killing NK cell. **(D)** Outcome of the 4-hour single-cell cytotoxicity assay in the microchip. NK:target cell ratio was 1: 4-7 (N=383). **(E-G)** Time-lapse sequence of NK cells killing either none (E), one (F) or three target cells (G) as acquired on the screening microscope. **(H)** Isolation of the identified serial-killer NK cell visualized by phase contrast (i, iv) and fluorescence imaging (ii, iii), as acquired on the cell-isolation microscope. **(i)** NK cell and target cells in the selected well prior to isolation. The NK cell is identified by its fluorescent signal (purple, arrowhead). **(ii)** Fluorescence image showing that the NK cell (arrowhead) has been lifted into the micropipette opening (dark circle). **(iii**) Fluorescence image showing that the NK cell (arrowhead) has been released into a different culture vessel for further analysis. **(iv)** Image showing that the NK cell was removed from its original well. Scale bars: 2 mm (B), 200 µm (C), 20 µm (E-H).

### Limitations

This study demonstrates the feasibility of combining single-cell functional assays in deep microwells and selective retrieval. Nevertheless, this approach has relatively low throughput, mainly due to slow cell isolation (Supp Fig 2), resulting in limited cell numbers (tens to possibly hundreds) for downstream investigations. Operating the instrument requires careful manual setup and supervision, a slight off-target approach into a well may result in the micropipette hitting the silicon wall and breaking. In that case, the piezo-actuated isolation device needs to be entirely dismounted and cleaned up, before restarting the entire setup and priming procedure. In practice, this aspect, along with potential clogging of the micropipette due to aspirating cell debris or bystander cells, further limits the effective throughput of the technique. The isolation throughput and success rate would most likely improve by using larger or more shallow wells. For this reason, we find that this approach should be reserved for specific investigations where the functional measurements are difficult to perform outside of deep microwells, and where low downstream cell numbers are acceptable.

## Discussion

In this study, we have implemented and evaluated a system for single-cell isolation from microwell arrays in a silicon-glass chip. The glass-bottom microwells are optimized both for high-content screening and high-resolution microscopy, and their deep and narrow, high-aspect ratio geometry makes them compatible with long-term cell analysis assays, even of highly motile suspension cells such as NK cells (28,44). This is in contrast to other microwell arrays that have been evaluated together with various cell isolation systems, that are either destructive (36–40) or rely on shallow wells often made in soft material (like PDMS) (23,24,31,41–43). The solution for single-cell retrieval that we evaluated involves a glass micropipette entering the wells from above and piezo-electric fluid actuation to pick and release the cell of interest. In our evaluation, we characterized the flow behavior of the system using numerical simulations and experiments with microbeads. We measured a 23 µm radius-of-effect for pipette aspiration in the microwells, which is thus the sorting resolution of our system when aiming at the center of the objects. Using beads, we found that picking two beads in close proximity to each other behaved as independent events, confirming that the aspiration of a first object did not negatively affect the fluid flow for aspiration of the second. Then we demonstrated single-cell isolation of primary NK cells with an average success rate over 60% and retained cell viability. We could successfully isolate single serial-killing NK cells despite them being surrounded by live and dead target cells. However, a risk related to aspirating bystander cells lies in the potential for contamination of downstream assays. For particularly sensitive assays and when the cells are not sufficiently well-separated, a two-step isolation process could be implemented, whereby the cells of interest are first transferred from their microwells to a larger vessel to remove possible contaminants, before a second round of isolation of the cells into their destination vessel (46).

With the workflow that we established during the evaluation, we demonstrated the potential of performing on-chip single-cell functional screening, followed by targeted isolation of specific cells based on the results from the screening assay. The high number of wells contained on the chip makes it possible to capture rare events (one in several thousand cells) and across four different biological conditions in parallel. Some high-content omics approaches can be performed directly on the isolated cells, such as proteomics by proximity extension assay (PEA) and single-cell RNA sequencing (47,48). Single cells obtained from isolation can be individually expanded into clonal populations for further functional analysis, or to generate cell counts compatible with other downstream assays such as flow or mass cytometry (49). It is also possible that such clonal populations could find direct use in cell-based immunotherapy.

From our evaluation, we conclude that the system is compatible with non-destructive, live cell isolation in deep, high-aspect ratio microwells made of silicon and glass. Specifically, it is possible to target individual cells in an 80 µm wide and 300 µm deep well containing multiple cells using piezo-actuated liquid aspiration with a thin glass micropipette. Likewise, it is also possible with the same system to dispense and release isolated cells into microwells. The current system may not be suitable for applications requiring isolation of thousands of cells, but it is feasible to isolate tens to possibly hundreds of individual cells from microwells efficiently within a reasonable experimental timeframe. This may be improved in the future by further automation and parallelization of the system.

## Materials and Methods

### Silicon-glass microwell chip

The silicon-glass microwell chips were produced by the SciLifeLab Customized Microfluidics facility according to a protocol previously described elsewhere (44,50). Briefly, a 300 µm-thick 4” silicon wafer was patterned through lithography and deep reactive ion etching, generating arrays with hexagonal-shaped through-holes. After oxidation, a 175 µm-thick glass wafer was anodically bonded to the bottom side of the micro-structured silicon wafer, thereby closing the holes and forming microwells with a flat glass bottom. The wafer was then diced into nine 22×22 mm^2^ microwell chips.

The polymer frame was made by mixing polydimethylsiloxane (PDMS) (Sylgard 184, Dow Corning) in a 10:1 ratio and molding it in a precision milled aluminum mold. After curing at 65°C for at least one hour, the polymerized frame was manually released from the mold and exposed to oxygen plasma together with the microwell chip for surface activation. Directly after, the two parts were aligned and put together to create a permanent bond. The chips were cleaned and reused for multiple experiments. The chip holder was made by precision milling of aluminum, which was black anodized after the fabrication.

### Culture of primary human NK cells

Human NK cells were isolated from blood from anonymous healthy donors, requiring no ethical permit according to local ethics regulations. Peripheral blood mononuclear cells (PBMCs) were separated from buffy coats by density gradient centrifugation (Ficoll Paque, GE Healthcare), and kept frozen at -152°C in Fetal Bovine Serum (FBS) + 5% dimethyl sulfoxide (DMSO) for storage. NK cells were then isolated from freshly thawed PBMCs by negative selection according to manufacturer’s instructions (Miltenyi Biotec). NK cells isolated using either protocol were maintained in IMDM medium containing 10% FBS, 1% penicillin-streptomycin, 1mM sodium pyruvate, and 1× non-essential amino acids (all from Sigma Aldrich), and activated with 100 U/mL IL-2. The cells were used within 48 hours after isolation.

### Isolation device priming

Bead and cell isolation was conducted using a piezo-actuated isolation device (CellSorter Scientific). The protocols for setup and initialization of the piezo-electric head were adapted from the manufacturer’s recommendations. Blunt-ended glass micropipettes with a 20 µm inner diameter at the tip (CellSorter Scientific) were coated with a siliconizing solution (Sigmacote, Sigma Aldrich), by dipping the glass tip into the solution and letting it fill a few millimeters by capillary forces. The coating solution was then manually expelled from the pipette using the priming module of the isolation device, before letting the coating dry for at least 8 hours.

The fluidic chamber of the isolation device was primed by manually aspirating desiccated MilliQ water and desiccated PBS through the micropipette, then closed with a drop of mineral oil (Sigma Aldrich) and the piezo-element was brought into contact with the 1×1 O-ring below.

The vertical and lateral alignment of the micropipette head and the microscope objective was performed following the manufacturer’s instructions (45 and personal communication with B. Szabó, CellSorter Scientific). In short, the micropipette was lowered over an empty dish, then slowly brought to contact with the bottom of the vessel. Contact was confirmed by exerting short lateral movements of the micropipette relative to the dish.

### Calibration

Because the closed liquid chamber is particularly sensitive to air bubbles that could remain trapped during priming, we confirmed the correct initialization of the isolation device before each experiment, by running a calibration round. For this purpose, 5 µm-wide silica beads (Bangs Laboratories) were seeded in a Petri dish (Corning) containing PBS, and 10 well-separated beads were randomly selected for isolation. The pipette was lowered 15 µm over the beads, and a -5 V voltage ramp was applied over 100 ms for picking (corresponding to an estimated volume of 1.1 nL (45)), and each bead was expelled again in the same dish by applying a voltage of +20 V ramping over 10 s (4.6 nL). A calibration with at least 5 beads successfully picked up and released was considered as passed. Otherwise, a new priming step and subsequent calibration was performed.

### Retrieval of beads and cells from microwells

Clean, sterile microchips were prepared by filling the chambers with MilliQ water and placing the chip in a desiccator, effectively removing any remaining air bubbles from the wells. The chambers were then filled with PBS (experiments with beads) or complete NK cell medium (experiments with NK cells) before seeding 5 µm silica beads (Bangs Laboratories) or human NK cells, stained with CellTrace Yellow (Thermo Fisher Scientific), at 2×10^4^ cells or beads per chamber. After 15 min seeding, beads or cells on the well walls were washed away by manual pipetting, and isolation medium at room temperature was added. For beads, PBS was used, and for NK cells, either complete L-15 or magnetic separation medium (MACS; PBS + 5% FBS + 2 mM EDTA). For picking, the micropipette was placed 15 µm over the object and a -2 V voltage ramp applied over 100 ms. Their subsequent releases were also performed in microwells, with a slower speed to keep the object close to the deposition site (+20 V voltage ramp over 10 s). After NK cell isolation, the cell culture medium in the deposition chamber was changed to complete NK cell medium with cytokines, and the cells were imaged by time-lapse microscopy for 18 hours to document survival and proliferation.

The success of individual retrieval events was classified by continuously following the process with wide-field microscopy and observing both picking and release. For distance measurements, images of the microchip tile were taken prior to isolation with a marking for the target positions of the micropipette, and after isolation. The distance from beads to the pipette center and to the well wall were measured in ImageJ (NIH).

### Integration in functional screening workflow

Single-cell cytotoxicity screening assays against K562 tumor targets were performed as previously described (28). Briefly, CellTrace Violet (Thermo Fisher)-labeled tumor target cells were seeded into the microchip primed with complete RPMI medium supplemented with SYTOX Green (Thermo Fisher), at which point a first screen of the microchip was acquired to assess target cell death at seeding. CellTrace Orange-labeled NK cells were then seeded into the wells, and cells still in suspension after 15 min were washed off before the imaging was started.

We have previously developed a custom MATLAB-based software for analysis of microchip screening datasets (28), where single cell multi-channel segmentation and mapping to microwells can be performed. This software was expanded to include the export of result files containing the cell coordinates, fluorescence intensities as well as interpreted well outcomes. Although initially designed for the interpretation of datasets from cytotoxicity assays, the variability of the parameters measured, and the flexibility of the output format make this software readily compatible with other functional readouts.

In this study, we have set up an interface bridging our image analysis program with the commercial software used to control the CellSorter isolation device. This interface is built in MATLAB (MathWorks) and communication with the CellSorter software is conducted using the import/export of compatible settings files. Scanning the corner tiles of the chip makes it possible to calculate the rotation of the chip relative to the microscope stage axis. The images from these corner positions are used to generate a map of all tiles and individual wells in the microchip. Results from a screening assay can then be imported in the interface and wells sorted according to the functional results. To specifically target individual cells rather than the well in its globality, we built-in cell detection in multicolor fluorescence images and mapping to the corresponding well address. The cell coordinates thus obtained are used to generate an isolation map in a format compatible with the CellSorter software.

### Flow profile simulations

Flow profiles were simulated using COMSOL Multiphysics 5.6. A three-dimensional representation of a pipette 15µm from the bottom of a hexagonal well or open dish was created. The flow was modeled using the Creeping Flow physics module, with the top of the pipette set as an outlet with a flow rate of 17.5mm/s (corresponding to the maximum flow rate induced by a -1V voltage ramp over 100ms) and the interface towards the liquid reservoir set as a free liquid inlet. All other surfaces (inner and outer surface of the pipette, well walls and bottom, dish bottom) were set as slip-free walls. A stationary flow solution was generated with a “Fine” finite elements mesh.

## Acknowledgements

We thank The Knut and Alice Wallenberg foundation (Grant No 2018.0106), The Swedish research council (Grant No 2019-04925), The Swedish foundation for strategic research (Grant No SBE13-0092), The Swedish childhood cancer foundation (Grant No MT2019-0022), The Swedish cancer foundation (Grant No 19 0540 Pj) for financial support.

## Author contribution

Q.V. designed the study, developed protocols, performed experiments, analyzed data and wrote the article. N.S. designed the study, co-supervised the work, developed protocols and wrote the article. H.v.O. designed the study, developed protocols and performed experiments. K.G. designed the study and performed experiments. K.O. analyzed data. T.F. developed protocols and performed experiments. B.Ö. designed the study, supervised the work and wrote the article.

## Supplementary material

**Supplementary movie 1:** Cytotoxicity assay in microchip, followed by identification and isolation of a serial-killing NK cell.

**Supplementary figure 1:**
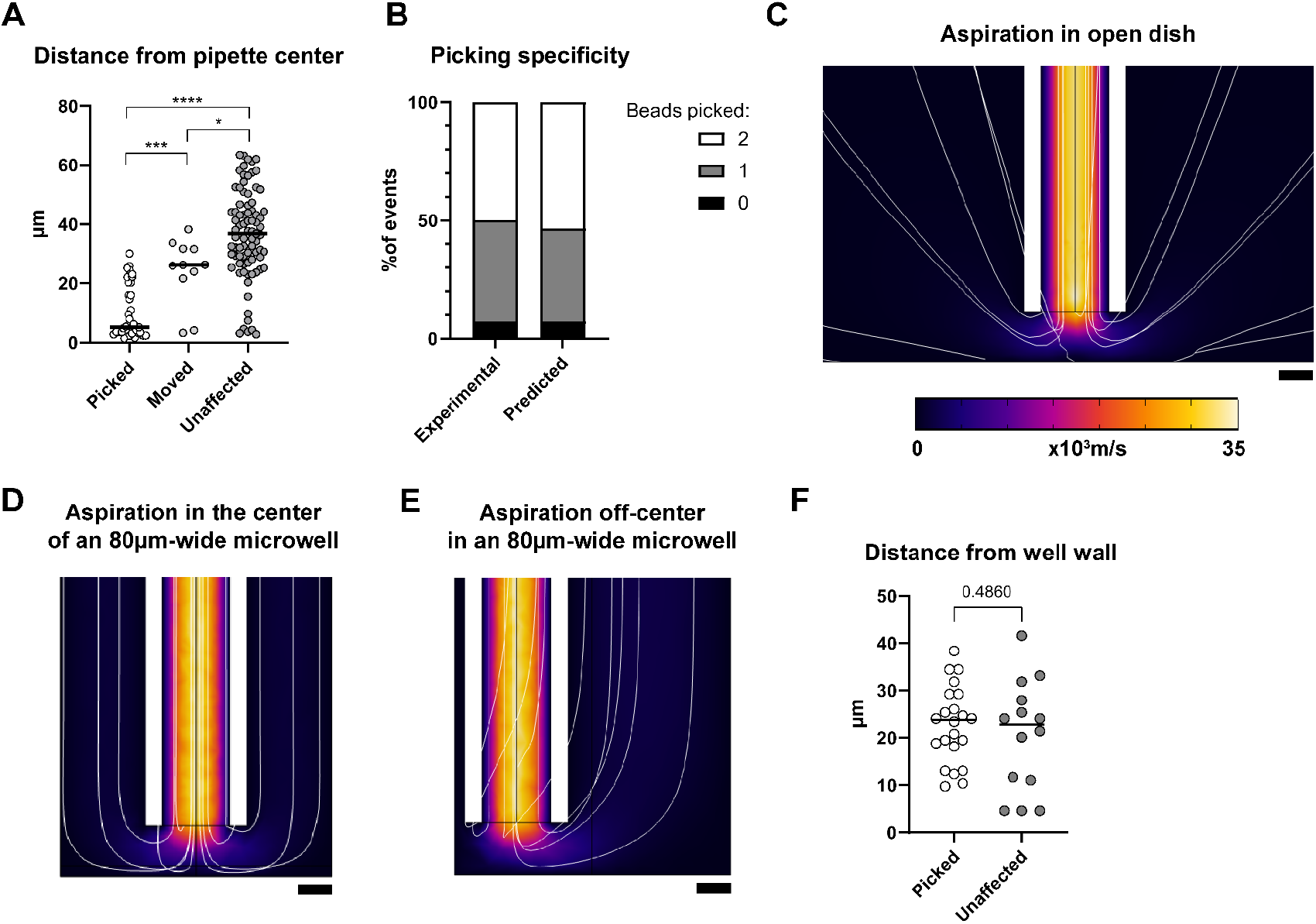
Characterization of the fluid flow profile around the micropipette tip. **(A)** During isolation of silica beads from the microwells, the effect of pipette aspiration on individual beads was documented and related to the distance from the bead to the center of the pipette. We identified 3 responses, either aspirated beads (“picked”), moved but not aspirated (“moved”) or unaffected. **(B)** For two beads lying within the area-of-effect of the pipette, we compared the outcome of picking of either or both beads (“Experimental”), to that predicted for independent events with an individual probability of 73% (“Predicted”). **(C-E)** Fluid flow profile during micropipette aspiration in an open culture dish (C), an 80 µm-wide hexagonal well with the pipette in the center (D) or off-center (E). Heat map: fluid flow velocity; white lines: velocity field lines; scale bars: 10 µm. **(F)** The effect of the proximity of beads to the well walls was investigated by considering a subset of picked and unaffected beads found within the area-of-effect of the pipette. The two groups (picked and unaffected) were chosen to have a matching average distance to the pipette center to compensate for any influence of that parameter. The average bead distance to the well wall was similar for the two groups indicating that the wall-effect was small.

**Supplementary figure 2:**
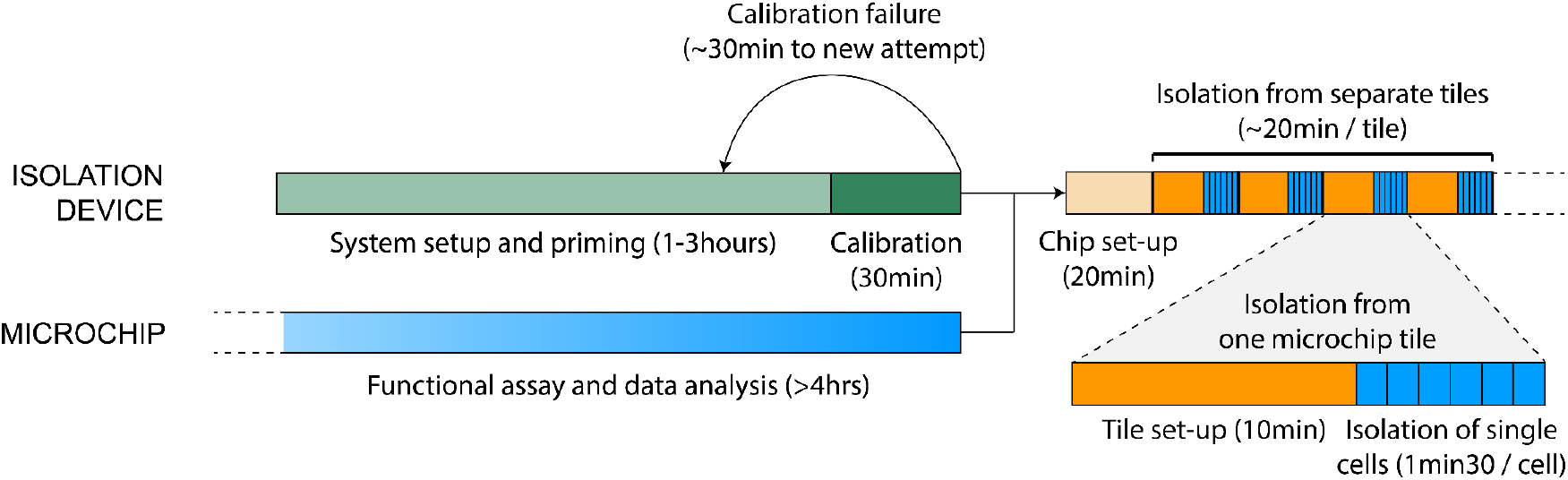
Time-course of cell isolation from the microwell chip. Setup of the isolation device can be started while functional experiments are being conducted on the microchip, since isolation does not necessarily have to be performed on the same microscope. Liquid priming of the device can be time-consuming as all air bubbles need to be expelled before closing the piezo-actuated chamber, and residual air bubbles would likely result in a failed calibration. A calibration is then conducted using silica beads in an open culture dish. Once the system is confirmed up and running, the chip can be installed for cell isolation. Chip setup is straightforward and simply requires scanning of corner positions of the microwell array to calculate all well coordinates, adjusted for chip rotation. Then, for every tile (sub-array of microwells) in the chip, a new image of the cells is captured and the cells of interest are marked out in a semi-automated fashion. Once setup is complete, the system automatically performs cell isolation, including targeting the cell, lowering the micropipette into the well, aspirating the cell, exiting the well and approaching the area selected for cell release, dispensing the cell, and refilling liquid in the micropipette at the designated reservoir.

***Supplementary movie 1: Functional screening and directed isolation of primary human NK cells***. *Primary NK cells were screened in the microwell chip for their cytotoxic potential against K562 tumor target cells. The example of a serial-killing NK cell is shown, which was automatically identified, then retrieved from its microwell with minimal effect on neighboring cells*.

